# Single Spectral Flow Cytometry Panel for Simultaneous Detection of Hematopoietic Stem, Progenitor, and Mature Lineages in Mouse Bone Marrow

**DOI:** 10.64898/2026.01.13.699036

**Authors:** Léa Girondier, Manon Richaud, Julien Grenier, Ana Belen Pérez-Oliva, Jenny Van Asbeck-van der Wijst, Michel Aurrand-Lions, Maria De Grandis, Christophe Lachaud, Cyril Fauriat, Laura Braud

## Abstract

Hematopoiesis occurs in the bone marrow of adult mammals and is supported by hematopoietic stem cells that sustain lifelong blood cell production. Pathological conditions can disrupt HSC differentiation, causing anaemia, immunodeficiency, and other cytopenias. Therefore, precise and simultaneous identification of hematopoietic populations from stem to mature cells is essential for understanding disease mechanisms and developing targeted therapies. In order to study these alterations, flow cytometry is generally a need-to-have monitoring technique. However, most cytometry panels are designed to thoroughly study a particular population, leaving out potential discoveries on other populations from the same microenvironment. Here we present the design of complex 19-marker spectral flow cytometry panel capable of simultaneously identifying HSPCs, erythroid, myeloid and lymphoid cells within a single murine BM sample. This integrated approach replaces multiple conventional panels and enables comprehensive mapping of hematopoietic differentiation from a single assay. Validation confirmed accurate detection of long-term and short-term HSCs, multipotent progenitors, common myeloid and lymphoid progenitors, and erythroid populations from proerythroblasts to reticulocytes. UMAP visualization captured the continuous trajectory of differentiation. We validated our method on aged and β-thalassemic mouse models showing a clear detection of hematopoiesis and erythropoiesis alterations respectively. This panel provides a robust, flexible, and scalable platform suitable for both basic research and translational studies.

## 1. Introduction

The bone marrow is the primary site of hematopoiesis in adult mammals, sustained by hematopoietic stem cells (HSCs) that generate all mature blood lineages throughout life. Pathological conditions can alter HSC differentiation, leading to cytopenias such as anemia or immunodeficiency. Therefore, the ability to precisely and simultaneously identify hematopoietic populations, from HSCs to fully differentiated cells, is essential to understand the mechanisms underlying these disorders and to develop targeted interventions.

Flow cytometry remains the reference standard technique to study hematopoiesis, providing high-resolution quantitative analysis of cellular diversity within the bone marrow. However, most conventional panels are designed to study a specific lineage, typically hematopoietic stem and progenitor cells (HSPCs) [1–3], erythroid populations [4,5], or myeloid and lymphoid compartments [6,7]. This strategy requires multiple samples, increases biological variability, and limits the integrated assessment of hematopoietic differentiation within a single biological sample.

To overcome these limitations, we developed and optimized a high-dimensional, single-sample spectral flow cytometry panel composed of 19 markers, enabling the comprehensive characterization of hematopoietic populations in a single murine bone marrow sample. This integrated panel simultaneously identifies HSPCs, myeloid, lymphoid and erythroid populations across all stages of maturation, from proerythroblasts to reticulocytes, within one assay.

Validation experiments confirmed accurate detection and discrimination of long-term and short-term HSCs (LT-HSCs, ST-HSCs), multipotent progenitors (MPP), common myeloid and lymphoid progenitors, as well as all erythroid maturation stages. UMAP-based visualization faithfully captured the continuous hierarchy of differentiation from stem to mature cells. Importantly, this single-tube, high-parameter approach consolidates three conventional panels into one, minimizes sample consumption, and allows reproducible mapping of hematopoietic trajectories within the same biological sample.

We further validated our approach under pathological conditions using aged mice known to display hematopoietic alterations, especially HSC dysfunction and a myeloid lineage bias [8]. We also used a mouse model of β-thalassemia (beta-thal), a congenital anemia caused by mutations in globin genes leading to defective erythropoiesis and chronic hemolytic anemia [9–11]. The panel successfully detected hematopoietic abnormalities previously described in these models, including progenitor expansion and erythroid maturation deficiency, confirming its sensitivity and applicability to pathological hematopoiesis.

Beyond a broad phenotypic characterization, our panel was explicitly designed to support the evaluation of emerging treatments by intentionally leaving specific fluorescence channels available for the detection of reporter proteins such as mCherry or GFP. As an example, therapeutic options for hematopoietic alterations, such as anemia remain limited, and many patients continue to experience suboptimal long-term outcomes. Recent advances in molecular and genetic therapies, including gene therapy and RNA-based approaches, rely on an expanding range of viral and non-viral delivery vectors to correct disease-causing defects at the level of hematopoietic stem and progenitor cells [12,13]. As these approaches are primarily developed and optimized in preclinical models prior to clinical translation, there is a critical need for robust and comprehensive tools to accurately evaluate hematopoietic progenitor targeting, as well as the global impact of these interventions on hematopoiesis and erythropoiesis. In this preclinical context, our single-tube, high-parameter cytometry panel provides a robust, flexible, and scalable platform ideally suited for early-stage therapeutic evaluation. By enabling the simultaneous characterization of all major hematopoietic compartments within a single assay, it allows (i) precise monitoring of hematopoietic progenitor targeting by therapeutic delivery vectors using reporter-based readouts, and (ii) an integrated assessment of treatment-induced alterations across the entire hematopoietic hierarchy.

This comprehensive and system-level view makes our panel particularly valuable for preclinical studies of next-generation therapies for hematopoietic disorders such as congenital anemias, and supports informed decision-making prior to clinical translation.

## 2. Materials

### 2.1. Reagents and equipment

All reagents and equipment used in this study are listed in Table 1.

**Table 1.** Reagents and Equipment.

| Product | Company | Reference |
| --- | --- | --- |
| Brilliant buffer | BD Bioscience | 566385 |
| True-Stain Monocyte Blocker | Biolegend | 426103 |
| Cell strainer 70 µM | Sigma | CLS431751 |
| Needle 21G | Terumo | AN*2138R1 |
| 16% formaldehyde (10x10ml) | Thermofisher | 28908 |
| Ultracomp eBeads | invitrogen | 01-3333-42 |
| Lysis buffer ACK | Gibco | A10492-01 |
| Equipment | Company |  |
| Spectral Cytometer | Cytek | - |
| Softwares | Company |  |
| OMIQ | GraphPad Inc. | - |
| GraphPad Prism | GraphPad Inc. | - |
| SpectroFlow | Cytek | - |
| CytekCloud (Panel Builder) | Cytek | - |

### 2.2. Antibodies

All antibodies used are listed in Table 2 including fluorochrome, supplier, clone, catalogue number and titration.

**Table 2.** Antibodies and titration.

| N° | Marker | Ch | Fluo | Host | Company | Cat# | Clone | Titration |
| --- | --- | --- | --- | --- | --- | --- | --- | --- |
| 1 | CD16/32 | UV2 | BUV395 | Rat | BD | 740217 | 2.4G2/50µg | 1:200 |
| 2 | CD71 | UV7 | BUV496 | Rat | BD | 750648 | OX-26 | 1:800 |
| 3 | F4/80 | UV11 | BUV661 | Rat | BD | 750643 | T45-2342 | 1:400 |
| 4 | CD3 | UV14 | BUV737 | Rat | BD | 612803 | 17A2 | 1:200 |
| 5 | CD45 | UV16 | BUV805 | Rat | BD | 568336 | 30-F11 | 1:400 |
| 6 | Sca-1 (Ly-6A/E) | V1 | BV421 | Rat | BD | 562729 | D7 | 1:800 |
| 7 | CD19 | V4 | BV480 | Rat | BD | 566167 | 1D3 | 1:800 |
| 8 | CD11b | V7 | Viogreen | Human | Miltenyi | 130-113-811 | REA592 | 1:400 |
| 9 | CD127 | V11 | BV650 | Rat | Biolegend | 135043 | A7R37 | 1:200 |
| 10 | CD150 | V13 | BV711 | Rat | BL | 115941 | TC15-12F12.2 | 1:400 |
| 11 | CD48 | B3 | RB545 | Hamster | BD | 756177 | HM48-1 | 1:400 |
| 12 | CD44 | B8 | PerCP | Rat | Biolegend | 103035 | IM7 | 1:200 |
| 13 | Ter119 | YG1 | PE | Rat | BL | 116207 | TER-119 | 1:800 |
| 14 | CD135 (Flt3) | YG9 | PE Vio770 | Human | Miltenyi | 130-111-153 | REA779 | 1:200 |
| 15 | CD4 | R2 | AF647 | Rat | BD | 557681 | RM4-5 | 1:800 |
| 16 | CD34 | R4 | AF700 | Rat | Invitrogen | 56-0341-82 | RAM34 | 1:100 |
| 17 | ViaDye Red | R6 | ViaDye Red | - | Cytek | R7-60008 | - | 1:50 000 |
| 18 | c-Kit (CD117) | R7 | APC Cy7 | Rat | BL | 105825 | 2B8 | 1:400 |
| 19 | Ly-6G/Ly-6C (Gr-1) | R8 | APC Fire 810 | Rat | BL | 108469 | RB6-8C5 | 1:400 |

### 2.3. Cytometer

The Cytek® Aurora cytometer was used in this study. Cytek® Aurora cytometer is configured with 5 lasers (355 nm (UV), 405 nm (Violet), 488 nm (Blue), 561 nm (Yellow-Green) and 640 nm (Red)) and a full-spectrum detection system comprising 64 avalanche photodiode detectors (UV1-UV16; V1-V16; B1-B14; YG1-YG10; R1-R8). Spectral reference controls and daily quality control (QC) were performed according to the manufacturer’s guidelines.

### 2.4. Software

Data analysis was performed in the SpectroFlow software (Cytek) and OMIQ (Licence software) and Cytek Cloud web tool. GraphPad prism was used to generate the graphs.

### 2.5. Animals

Mice Wild-Type (WT) (young and aged) and beta-thalassemic (beta-thal) were housed under controlled conditions of temperature (21±1°C), hygrometry (60±10%) and lighting (12 h per day). Animals were acclimatized in the laboratory for one week before the start of the experiments. All animals received care according to institutional guidelines, and all experiments were approved by the Institutional Ethics committee number 16, Paris, France (beta-thal mice: strain B6.D2-Hbbd3th/BrkJ, licence number #49620-2024052208518465). Mice were euthanatized and organs and blood were collected and processed for further evaluations.

### 2.6. Statistical analysis

Data are expressed as mean values ± standard error of the mean (SEM). Statistical significance was tested using either one or two-way analysis of variance (ANOVA) with Fisher multiple comparison test. The results were considered significant if the p-value was <0.05.

## 3. Methods and Results

### 3.1. Objective and Panel design

This panel was primarily designed in order to precisely identify hematopoietic stem cells (long-term LT-HSC, short-term ST-HSC, multipotent progenitor MPP2, MPP3, MPP4), progenitor subsets (common myeloid progenitor CMP, megakaryocyte erythroid progenitor MEP, granulocyte-monocyte progenitor GMP and common lymphoid progenitor CLP) and erythroid populations (from pro-erythroblasts to reticulocytes) using a single high-dimensional spectral flow cytometry panel (Figure 1). In addition, the panel allowed a broader identification, though with less resolution, of mature myeloid cells, granulocytes, dendritic cells, macrophages, and lymphoid cells, (B and T cells). The stepwise workflow for spectral flow cytometry experiment is illustrated below and in Figure 1.

**Figure 1.**
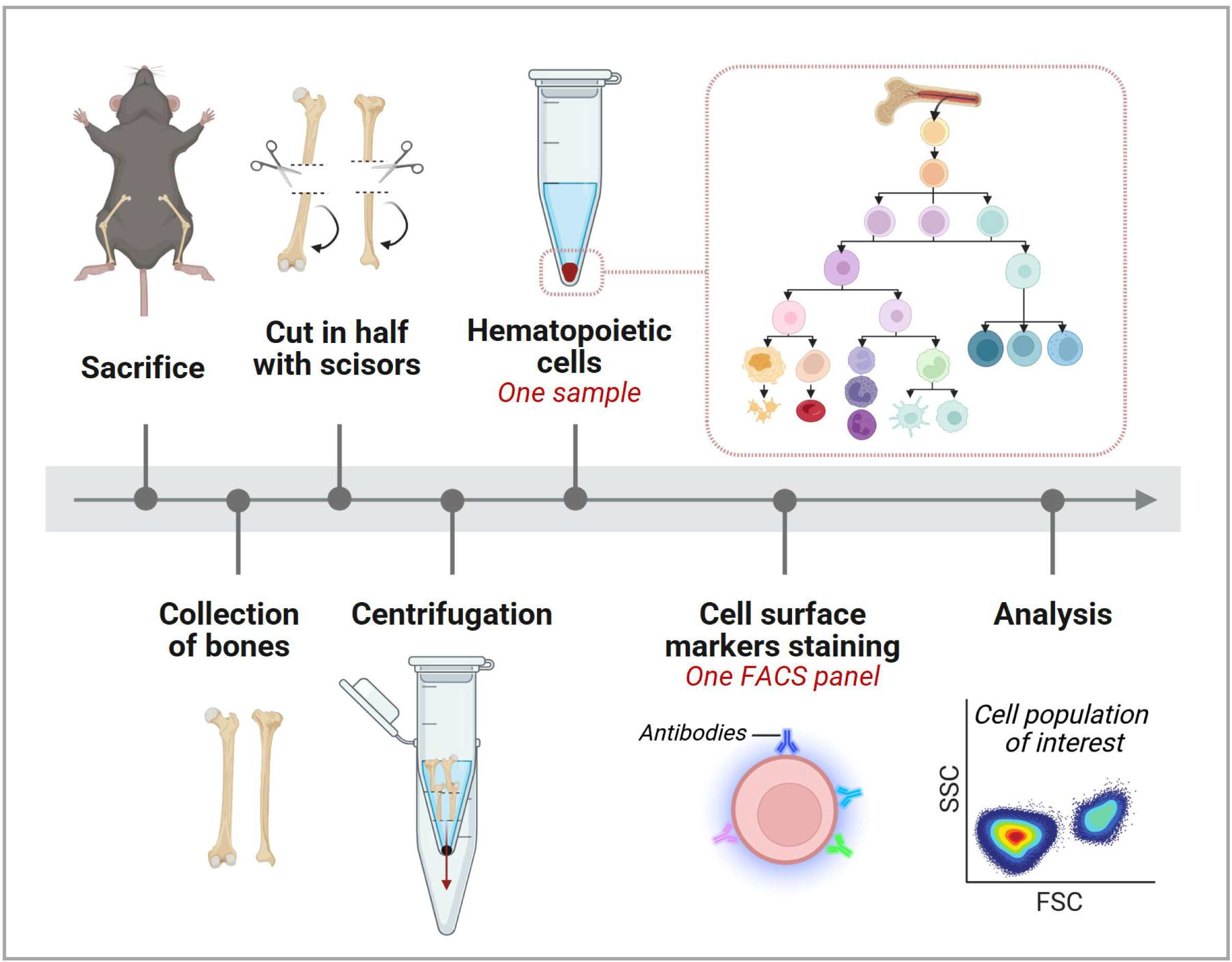
Schematic representation of the experimental procedure. *LT-HSC: Long-term hematopoietic stem cell; ST-HSC: Short term HSC; MPP: multipotent progenitor; CMP: common myeloid progenitor; GMP: granulocyte-monocyte progenitor; MEP: megakaryocyte erythroid progenitor; Pro-E, Baso-E, Poly-E, ortho-E: pro, basophilic, polychromatic, orthochromatic erythroblasts*

#### 3.1.1. Defining bone marrow cells to analyze

The initial step in panel development consists of defining the relevant cell populations within the biological sample, in this case, murine bone marrow, and identifying the most informative surface markers for their discrimination. Based on this analysis, we identified and selected 19 markers to distinguish 19 cell types of the mouse bone marrow (Figure 2A-B). The scientific references and validation for the selected cell-surface markers are provided in Section 3.3.(Data Analysis).

**Figure 2.**
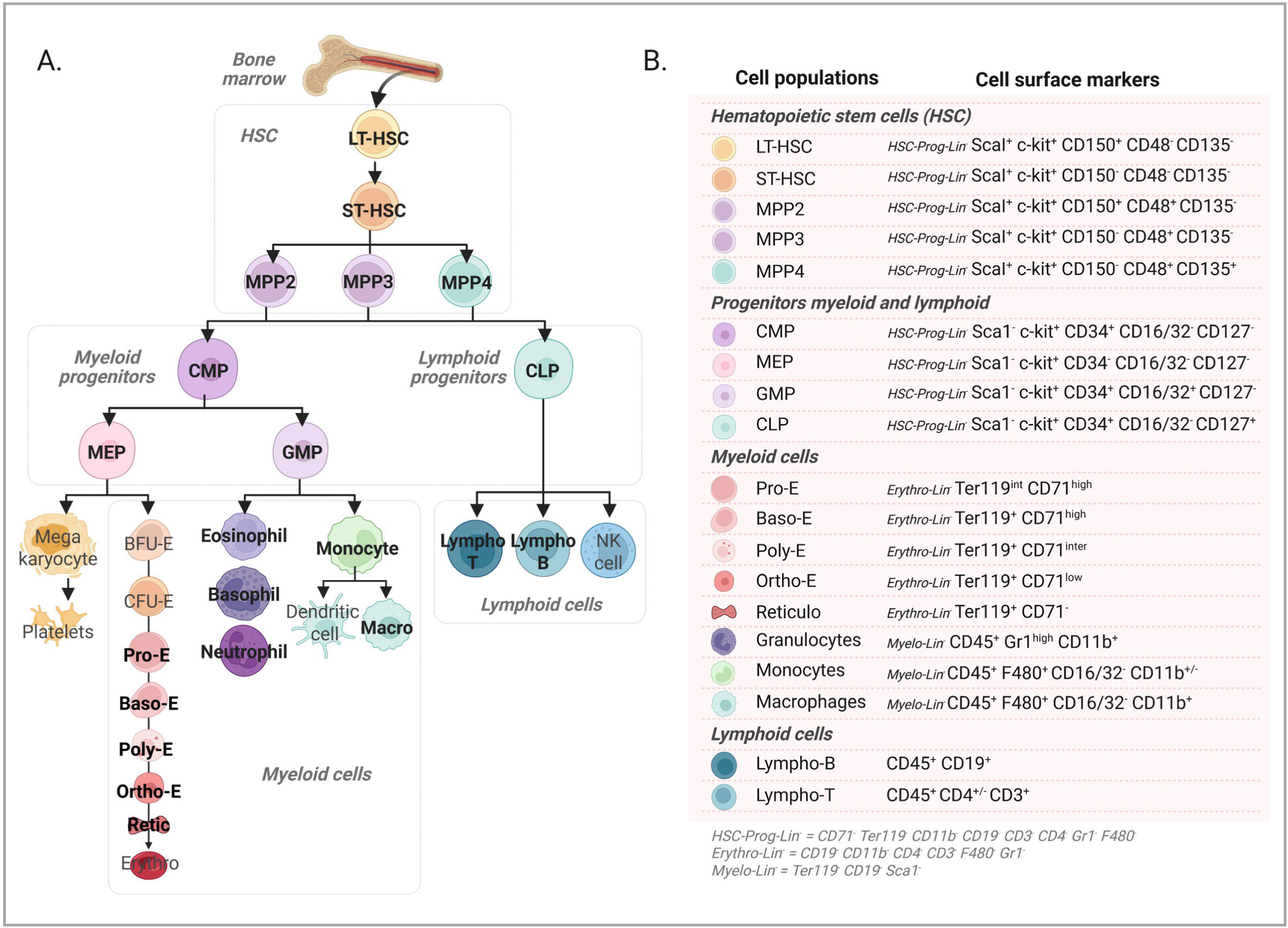
Cell surface markers of bone marrow cell populations. (A) The schema pictures a simplified view of the hematopoietic system, with a focus on the compartment that can be analyzed with the proposed flow cytometry panel. (B) Details of the main surface markers used for cell annotation in bone marrow sample.

#### 3.1.2. Panel design

We then designed the panel (Figure 3A) by selecting the most appropriate fluorochrome for each antibody based on several criteria. First, fluorochromes were evaluated according to their full emission spectra (Figure 3B) and their similarity index profiles (Figure 3C), using the Cytek Cloud online tool to minimize spectral overlap. The similarity index was used to assess spectral proximity between fluorochromes, and combinations with a similarity index above >0.85 were avoided to prevent unmixing errors and loss of resolution. Additional considerations included fluorochrome brightness and antigen expression pattern, low-fluorescence for high-expressed markers, and commercial availability of the required antibody (Table 2).

**Figure 3.**
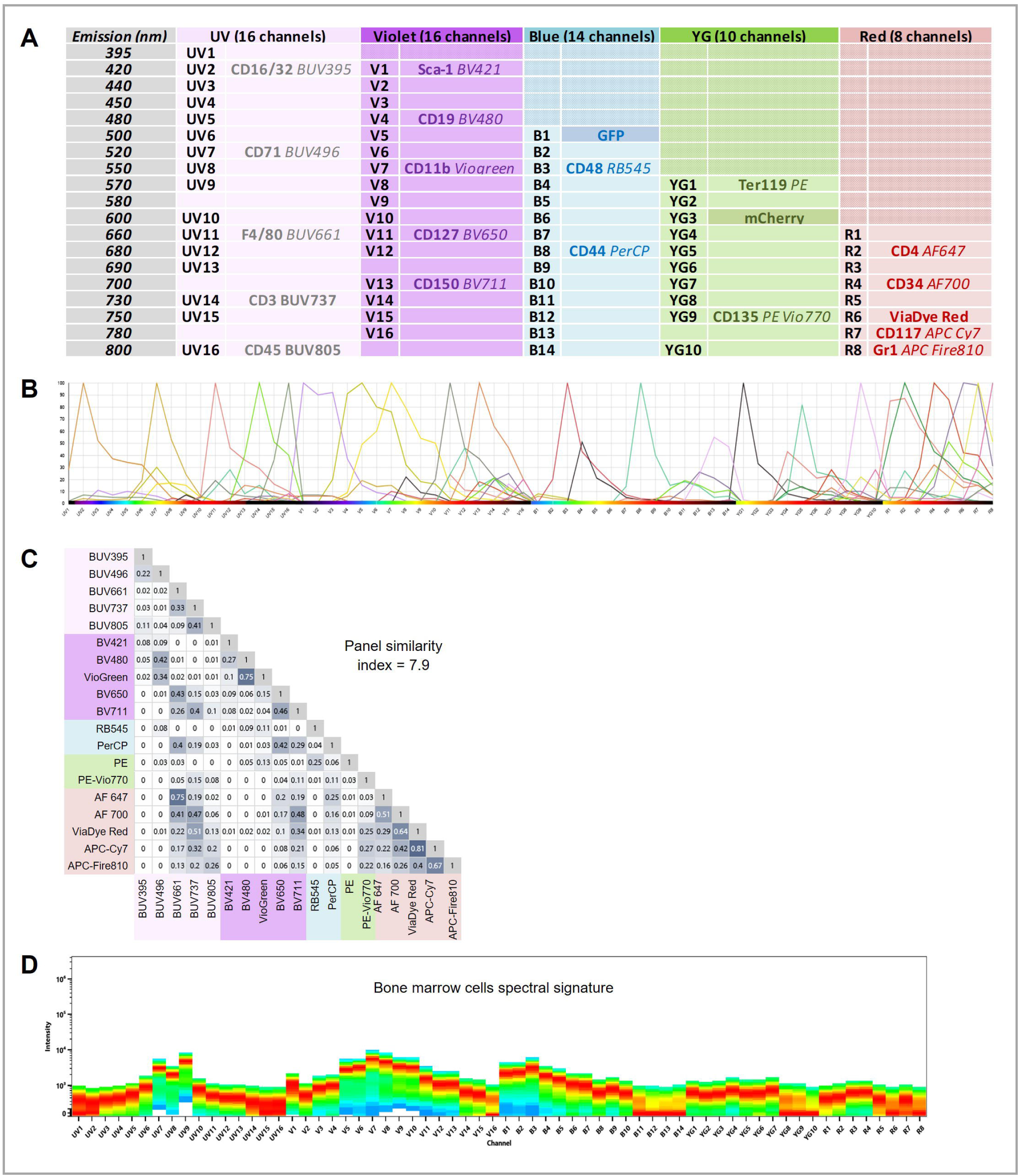
Spectral flow cytometry panel design. (A) The spectral cytometer used in this study is a Cytek Aurora that analyses the signal through 64 diode avalanche detectors. The principal emission peak of each fluorochrome is indicated together with the associated antibody. The list of antibodies used in the study is presented in Table 2. (B) Spectral signature of the different antibodies selected. (C) The graph depicts the index similarity (spectral overlap) of each fluorochrome and the overall similarity index of the panel (Cytek web tool). (D) The spectral signature of the unstained bone marrow cells (background) which is deduced from the other fluorochrome specific signals.

During panel design, we also characterized the intrinsic autofluorescence of bone marrow cells to guide fluorochrome selection (Figure 3D). By identifying the channels with the highest autofluorescence (V7 and B3), we strategically assigned well expressed antigens to these channels, ensuring optimal signal detection while minimizing interference from the background autofluorescence.

#### 3.1.3. Antibody Titration and Optimization

Once the panel has been designed, all antibodies were titrated to determine their optimal concentration in order to achieve an optimal balance between signal level for positive populations and negative populations (Table 2). Titration aimed to achieve maximal resolution between positive and negative populations while minimizing background signal and spillover. If a selected fluorochrome was not optimal, either due to overlap with another fluorochrome leading to excessive spectral spillover, insufficient signal intensity or limited availability, the panel was adjusted accordingly, replacing the fluorochrome or redistributing markers to achieve an optimal panel design. The overall similarity index for our 19-fluorochrome panel was 7.9 (Figure 3C).

### 3.2. Step-by-step experimental procedure

#### Step 1. Isolation of Bone Marrow Cells

Tibia and femur were collected in PBS 1x and divided into fragments, each bone was cut in half and placed with the cut surface down into a pierced tube nested within an unpierced tube (pierced tubes were prepared using a heated 21G needle) [14] (Figure 1). 500 µL of PBS 1x was added per tube. Samples were centrifuged at 3500g for 5 min to flush bone marrow into the lower tube, and the supernatant was removed. Pellets were resuspended in 1 mL PBS 1x, filtered through a 100-µm strainer, centrifuged at 400g for 5 min, and resuspended again in PBS. An aliquot of cell suspension was taken and incubated with RBC lysis buffer before counting. The equivalent of 5 million cells from the non-lysed fraction was transferred into 35-µm filter-cap FACS tubes to obtain cell suspensions for staining (Figure 1). Of note, if not interested in erythroid cell measurement, one may choose to remove red blood cells by using RBC lysis buffer on the whole bone marrow cell suspension.

#### Step 2. Preparation and Staining of Cells

##### Viability Staining

Cells were first washed once in PBS in order to removed excess of soluble proteins in the supernatant, then incubated for 20 minutes with 100µL of the viability dye ViaDyeRed mix, prepared at a 1:50,000 dilution in PBS. Following incubation, cells were washed with PBS, and subsequent staining steps were performed as described below. Antibody dilutions are listed in Table 2.

##### Pre-incubation with CD34 and CD127 antibodies

Cells were initially incubated with CD34 in a total volume of 20 µL for 5 minutes at room temperature. Then, without any intermediate wash, an additional 20 µL containing CD127 was added, and the cells were incubated for 15 additional minutes at room temperature. This two-step pre-incubation with CD34 and CD127 insured a better signal during acquisition.

##### Surface Staining – Mix 1 (Mix Blue–YG–Red)

Without any prior washing, cells were sequentially stained with two antibody mixes. First, 30 µL of the Blue–YG–Red Mix 1 (2× concentrated) was added to each sample. This mix contained antibodies against CD44, CD48, Ter119, CD135/Flt3, CD4, c-Kit/CD117, and Gr-1, as well as the TrueStain Monocyte Blocker to reduce nonspecific fluorochrome-cells interaction such as Cy7 with myeloid cells.

##### Surface Staining – Mix 2 (Mix BUV–V)

Immediately after, 30 µL of the BUV–V mix 2 (2× concentrated) was added. This second mix included antibodies for CD16/32/FcyR, CD71, F4/80, CD3, CD45, Sca-1, CD19, CD11b, and CD150. Brilliant Stain Buffer was incorporated into this staining step to minimize fluorochrome interactions, following the manufacturer’s recommendations.

##### Final incubation and fixation

Samples were incubated for 20 minutes at room temperature in the dark, washed to remove excess antibodies, and finally resuspended in PBS containing 2% paraformaldehyde to fix the cells prior to flow cytometry acquisition.

#### Step 3. Data Acquisition

Data were acquired on a Cytek Aurora Spectral Flow Cytometer (Cytek Biosciences) equipped with 5 lasers ultraviolet, violet, blue, yellow-green, and red, allowing for high-dimensional spectral analysis. The instrument’s Cytek assay settings were configured to ensure optimal resolution of cell populations. Forward Scatter Area (FSC-A), which correlates with cell size, allowing clear discrimination between small debris and intact cells. Side Scatter Area (SSC-A), reflecting cellular granularity ensure proper cell doublets discrimination in combination with FSC signals. A signal threshold of 50,000 was applied to exclude sub-threshold particles and debris from acquisition.

### 3.3. Data Analysis

#### 3.3.1. Spectral Unmixing and Compensation

Spectral unmixing and fluorescence compensation were performed using the SpectroFlo software (Cytek Biosciences). For the unmixing process, the cellular signature of the unstained control mouse bone marrow sample was used as the reference to accurately capture and extract the sample’s intrinsic autofluorescence (Figure 3D). This autofluorescence was then computationally subtracted by the software [15]. Compensation beads (Invitrogen 01-3333-42) were included as reference controls for each antibody used in the panel.

Following unmixing, fluorescence compensation was applied, requiring only minimal adjustments (Suppl Fig 1). Once unmixing was completed, the unmixed gated “all events” were exported as FCS files and uploaded into the OMIQ software for further analysis.

#### 3.3.2. Manual Gating strategy using OMIQ software

Downstream analysis and identification of cell populations were carried out using OMIQ software. Standard gating strategies (FSC-A vs SSC-A and viability dye vs SSC-A) were applied to exclude doublets and dead cells, followed by manual (supervised) population identification of hematopoietic stem and progenitor cells (HSPCs), erythroid, myeloid and lymphoid mature cells (Figure 4, Figure 5 and Figure 6 respectively).

**Figure 4.**
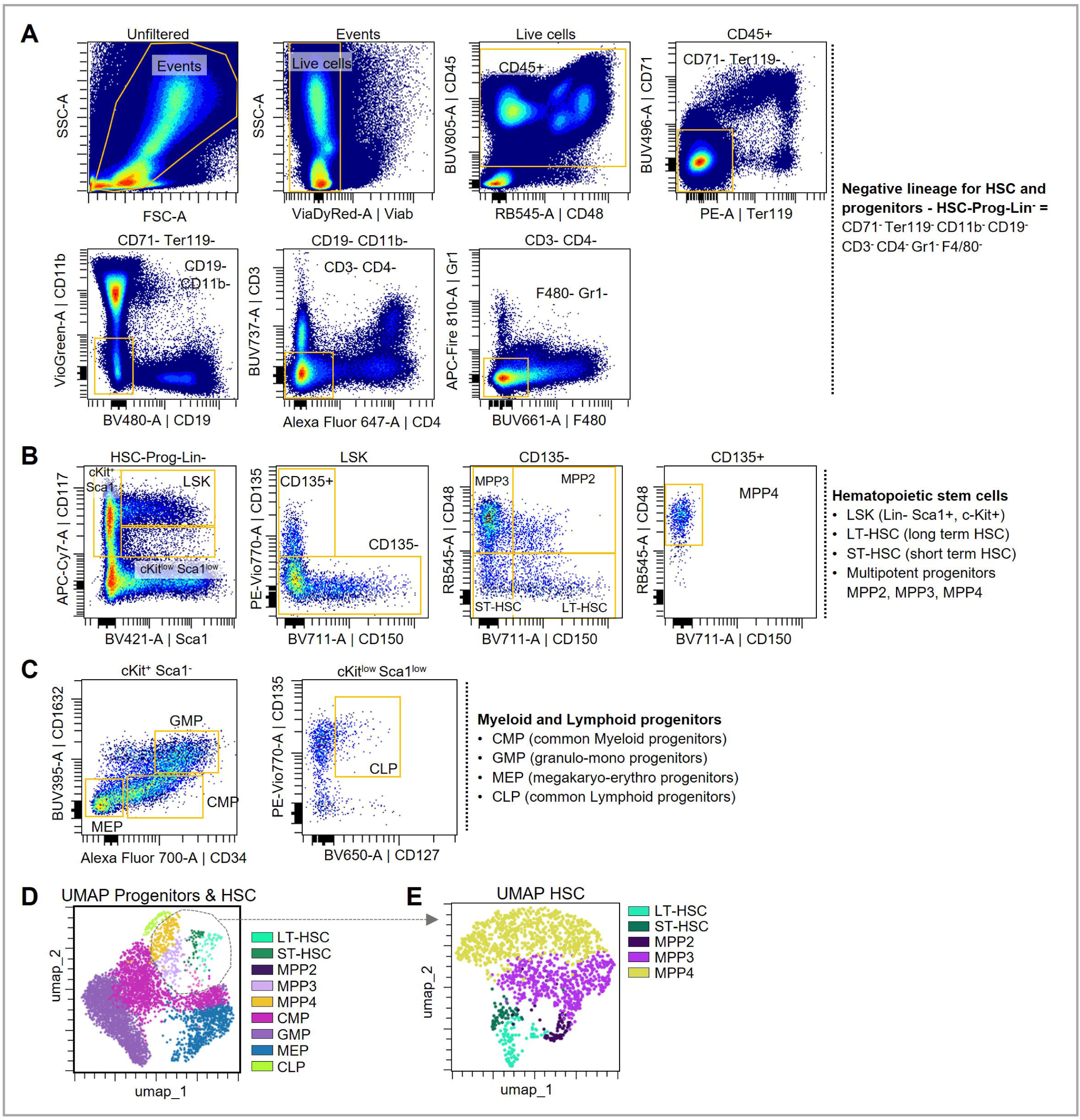
Identification of HSC and Progenitors. (A) Gating strategy for lineage negative for HSC and progenitors. (B) Gating strategy for HSC. (C) Gating strategy for Progenitors. (D) Unsupervised analyses (UMAP) for HSC/progenitors based on the expression of CD16/32, Sca1, CD150, CD127, CD48, CD135, CD34 and CD117 or (E) only HSC based on the expression of Sca1, CD150, CD127, CD48, CD135 and CD34. The indicated populations annotations originate from a manual gating superimposed on the UMAP representations.

**Figure 5.**
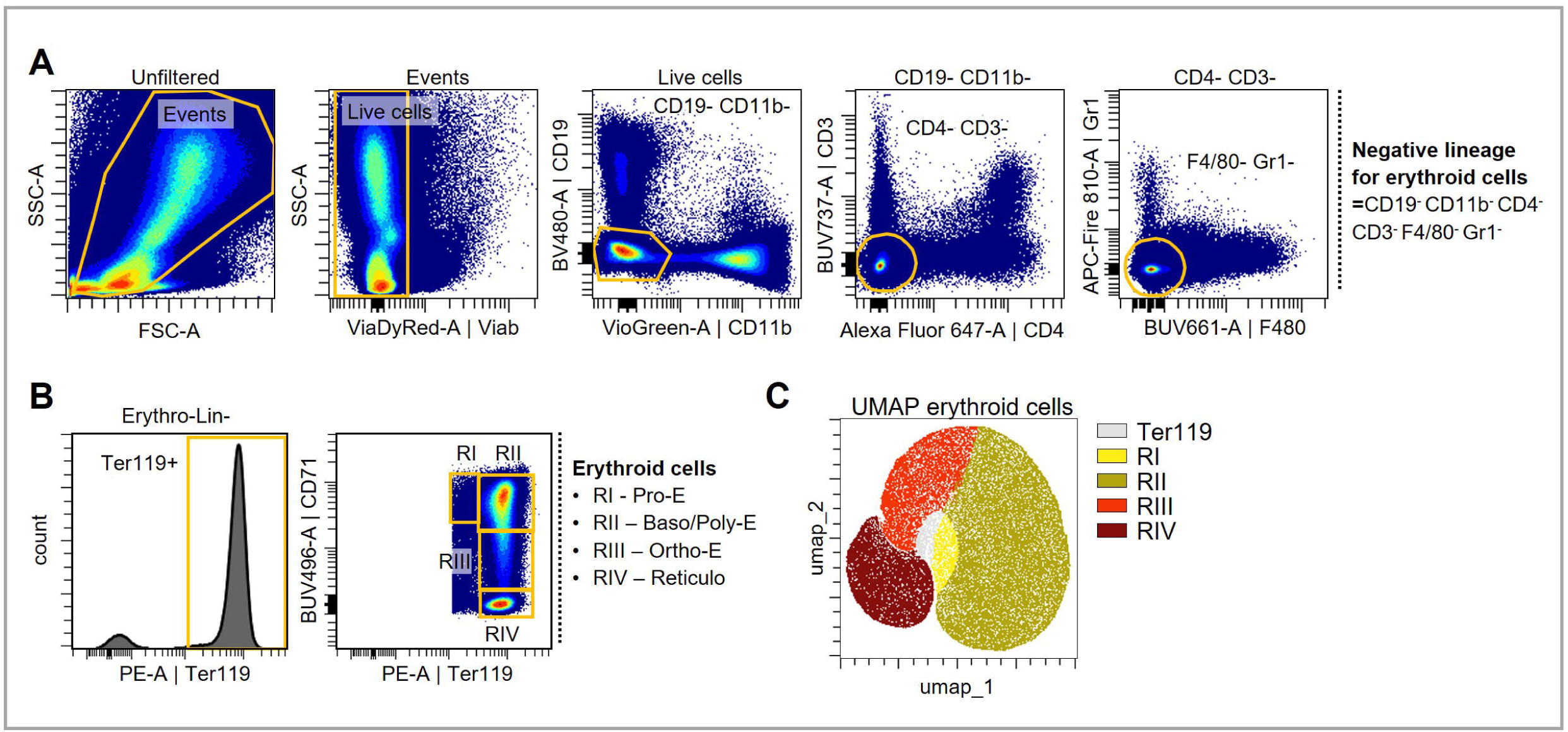
Identification of erythroid cells. (A) Gating strategy for lineage negative for erythroid cells. (B) Subsequent gating strategy for erythroid cells. (C) UMAP representing the organization of erythroid lineage based on the expression of CD71 and Ter119.

**Figure 6.**
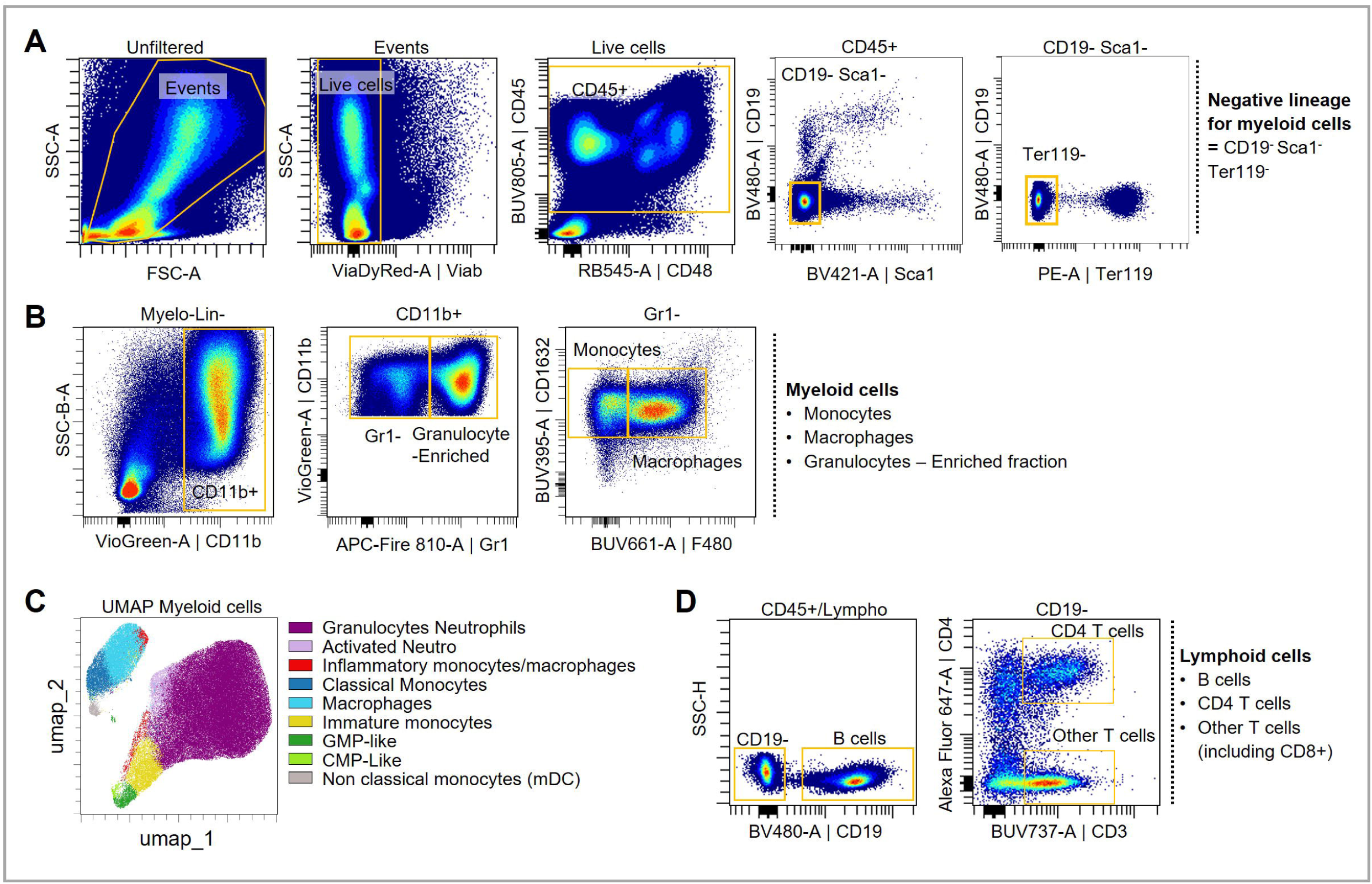
Identification of mature immune cells. (A) Gating strategy of lineage negative for myeloid cells. (B) Gating strategy of monocytes, macrophages and granulocytes. (C) UMAP of myeloid cells based on the expression of CD16/32, F480, CD11b, CD34, CD117 and Gr1. (D) Gating strategy for T and B cells

##### Quality controls

Prior to further analysis, we recommend to perform different quality controls. Signals may fluctuate during the time of sample acquisition, generating erroneous events. These events, which include clogs, flow speed, etc…, may influence data analyses at various range such as generating “new populations” (worst and rarer cases), altering fluorescence intensity, increase sample variability. Here, we have used the PeacoQC algorithm available in OmiQ [16]. This algorithm will remove events presenting abnormal signals in any of the parameters during time of acquisition, for each sample, generating a more reliable dataset (Suppl Fig 2).

##### Hematopoietic stem and progenitor cells (HSPCs)

Hematopoietic stem cells (HSCs) and progenitors were isolated using a lineage-negative (called here HSC-Prog-Lin^−^) pre-gating strategy to exclude mature hematopoietic populations (Figure 4A). Cells expressing CD71/Tfrc, Ter119, CD11b, CD19, CD3, CD4, Gr1, or F4/80 were excluded, thereby removing erythroid, myeloid, lymphoid differentiated lineages from the analysis.

Within the HSC-Prog-Lin^−^ fraction, the HSPC subsets were further defined based on expression of Sca1, c-Kit (CD117), CD150, CD48, and CD135, as previously described [1,2]. Using these markers, long-term HSCs (LT-HSCs), short-term HSCs (ST-HSCs), and multipotent progenitors (MPP2, MPP3, MPP4) were distinguished, enabling precise separation of self-renewing stem cells from more differentiated progenitor populations (Figure 4B).

Progenitors engaged in myeloid and lymphoid lineages were identified based on expression of CD34, CD127, CD16/32 and c-Kit (CD117) within the HSC-Prog-Lin^−^ fraction, as previously described [3]. This allowed the identification of the common myeloid progenitors (CMP), myelo-erythroid progenitors (MEP), granulocyte-monocyte progenitors (GMP) and common lymphoid progenitor (CLP) (Figure 4C).

For high-dimensional, unsupervised analysis, a random subsampling of 2,700 LSK (Lin^−^, Sca1^+^, c-Kit^+^) and a random subsampling of 14,000 c-Kit^+^, Sca1^+^ cells were performed and were used to generate a Uniform Manifold Approximation and Projection (UMAP) representation of HSPCs (Figure 4D-E). Antibody expression levels were visualized by using color-continuous visualization, which allowed assessment of the relative fluorescence intensity for each marker across cell populations. The fluorochrome associated with each antibody was mapped onto the Z-axis, providing a three-dimensional view of marker expression and ensuring accurate identification of each HSPC subsets (Suppl Fig 3A-B).

##### Erythroid cells

Erythroid cells were labeled and gated based on surface expression of Ter119 and CD71, as previously described [5,17]. Prior to erythroid-specific gating, a lineage-negative for erythroid cells (Erythro-Lin^−^) pre-gating strategy was applied to exclude non-erythroid cells. Cells positive for CD19, CD11b, CD4, CD3, F4/80 or Gr1 were excluded, thereby removing B cells, T cells, monocytes/macrophages, and granulocytes from the analysis (Figure 5A). Within the Erythro-Lin^−^population, Ter119-positive cells were then identified as erythroid lineage (Figure 5B). Within this population, sequential analysis of CD71 expression allowed discrimination of distinct stages of terminal erythroid differentiation: the morphologic characteristics broadly corresponded to pro-erythroblasts in the Ter119^int^ CD71^high^ cell population (Figure 5B, region I), basophilic erythroblasts in the Ter119^high^ CD71^high^ cell population (Figure 5B, region II), late basophilic and polychromatophilic erythroblasts in the Ter119^high^ CD71^int^ cell population (Figure 5B, region III), and orthochromatophilic erythroblasts/reticulocytes in the Ter119^high^ CD71^low^ cell population (Figure 5B, region IV) [5,17]. Cell surface marker expression levels were validated using a color-continuous visualization for each marker across the cell populations (Suppl Fig 4). We then performed a subsampling of 50,000 Ter119⁺ cells to generate a UMAP representation of the erythroid populations, allowing visualization of the continuous progression of erythroid maturation (Figure 5C).

##### Myeloid and Lymphoid mature cells

Finally, myeloid and lymphoid cells were labeled and gated based on specific surface markers, as previously described [6,7]. Prior to subset-specific gating, a myelo-lineage-negative pre-gating strategy (Myelo-Lin^−^ = CD19^−^, Sca1^−^, Ter119^−^) was applied to exclude erythroid and stem/progenitor cells (Figure 6A). Within the Myelo-Lin^−^ population, a CD11b positive gated was drawn to focus on the myeloid lineage. The granulocytes-enriched fraction was then defined based on Gr-1 expression, and monocytes and macrophages were distinguished using F4/80 and CD16/32 (Figure 6B). Notably, the use of Gr-1 antibody which targets both Ly-6C and Ly-6G antigen may preclude a clear distinction between the prominent population of neutrophils (Ly-6C-Ly-6G+) and inflammatory monocytes (Ly-6C+ Ly-6G-) which are less abundant. A non-supervised analysis (Figure 6C), using CD11b, Gr-1, CD16/32, F4/80 markers may refine the cell-type definition, although one should also consider adding markers such as Ly-6C, Ly-6G, SIGLEC-7, CD115 to better define the myeloid cell content. Cell surface marker expression levels were validated using a color-continuous visualization for each marker across the myeloid populations, which could reinforce the tentative cell type annotation that we performed with the UMAP (Suppl Fig 5). Lymphocytes were identified using CD19, CD3 and CD4 cell surface marker (Figure 6D). The use of markers such as CD1d, NK1.1, or anti-TCRγδ, would allow a better distinction of lymphoid subsets, with the risk of panel’s similarity index increase and loss in the quality of the panel.

##### Observation of all hematopoietic cells

In order to fully picture the content of the bone marrow hematopoietic system, we generated an UMAP integrating all identified hematopoietic cells (Figure 7). Cell surface marker expression levels were validated using a color-continuous visualization for each marker across the all-cell populations (Suppl Fig 6). This visualization clearly showed the most immature cells at the center of the UMAP and more mature cells (Lymphoid, myeloid and erythroid lineages) radiating in separate directions.

**Figure 7.**
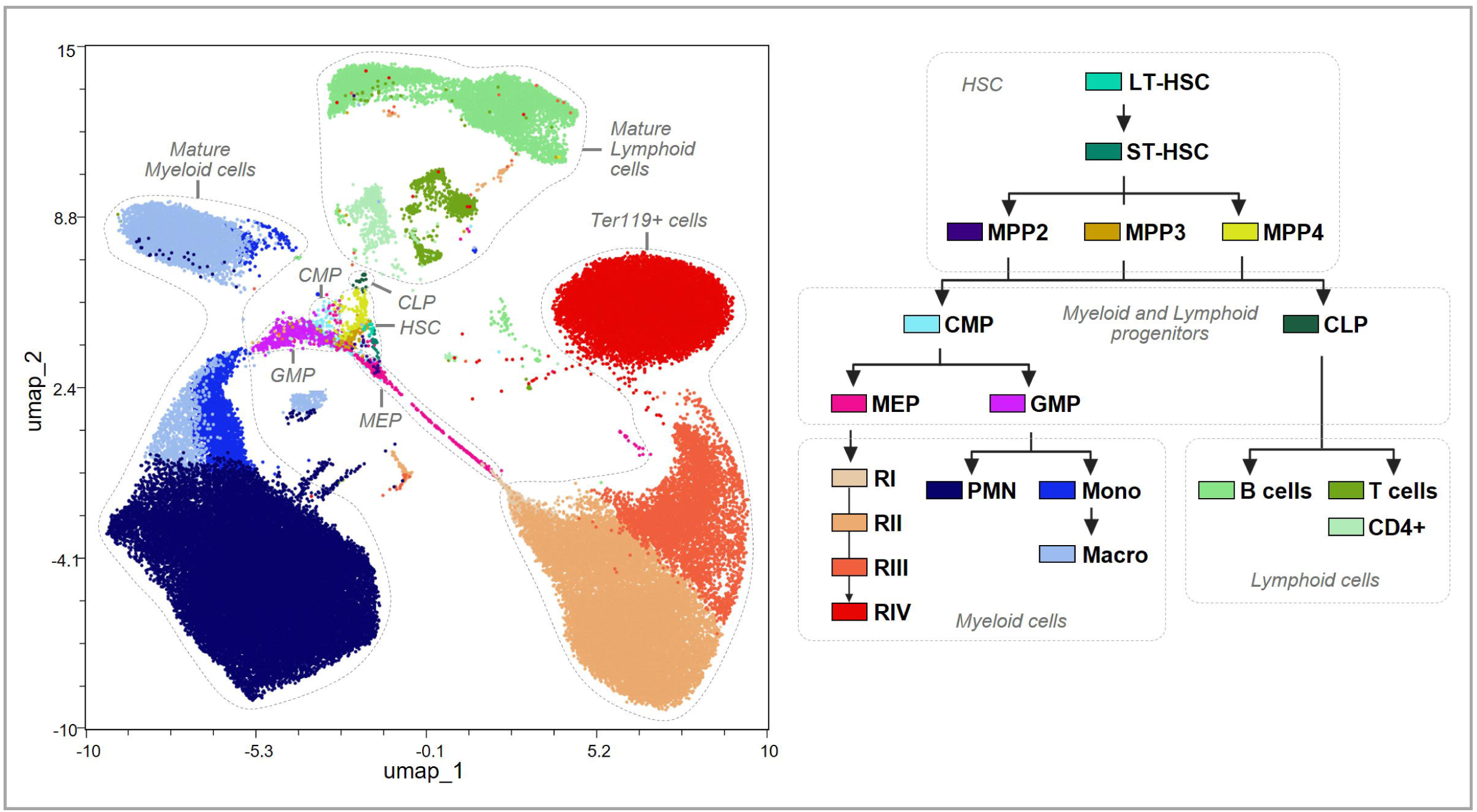
UMAP of all cells. UMAP analysis (left panel) was performed on 100,000 CD45^+^ Live cells using all parameters available in the panels. The cell annotation was superimposed after the manual gating presented in previous figures and the color code is presented on the right panel. *LT-HSC: Long-term hematopoietic stem cell; ST-HSC: Short term HSC; MPP: multipotent progenitor; CMP: common myeloid progenitor; GMP: granulocyte-monocyte progenitor; MEP: megakaryocyte erythroid progenitor; RI: pro-erythroblast; RII: basophilic erythroblast; RIII: polychromatic erythroblast; RIV: orthochromatic erythroblast; PMN: polymorphonuclear cells*

### 3.4. Panel validation

We validated our spectral cytometry panel on well-characterized mouse models known to exhibit hematopoietic alterations, in order to assess its ability to resolve changes in hematopoietic heterogeneity under pathological conditions.

First, we analyzed bone marrow cells of young and aged mice to confirm the panel capacity to detect age-associated hematopoietic alterations, as aging is known to affect HSC function and to drive a myeloid lineage bias [8]. Consistently, our analysis revealed clear age-related changes: aged mice (18 months) exhibited a significant expansion of LT-HSCs alongside a reduction in MPP4 populations (Figure 8A). These observations are in line with the myeloid bias that emerges during hematopoietic aging, reflecting both the shift in lineage commitment in mice and comparable trends reported in healthy elderly humans [18].

**Figure 8.**
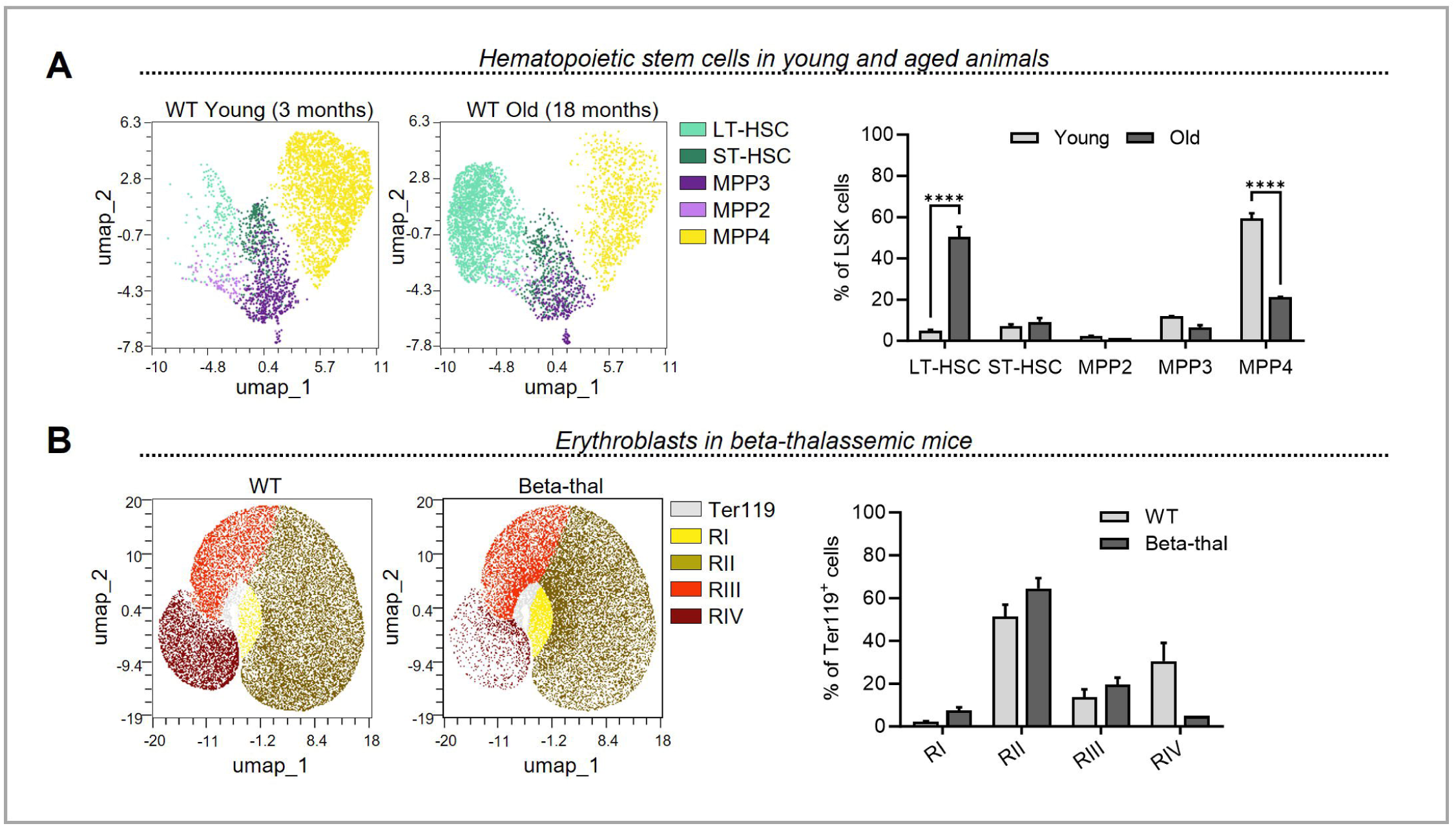
Application on different mouse models. (A) Analysis of HSC and progenitor cell frequencies in the bone marrow of young (3 months) and old (18 months) mice. The unsupervised analysis (UMAP on the left panel) clearly shows a discrepancy between cell density. The quantification of cell populations (right panel) confirms the increase in LT-HSC frequency and the loss of lymphoid-biased progenitors MPP4 (n=2/group). (B) UMAP analysis on erythroid lineage (left panel) showing the loss of the more mature erythroid cells (RIV) in mice suffering from beta-thalassemia (n=2/group). Quantification of cell frequencies confirms the loss of RIV in the beta-thalassemia group (right panel).

Next, we applied our panel to a beta-thalassemic mouse model (B6.D2-Hbbd3th/BrkJ), which is characterized by severe erythroid defects, ineffective erythropoiesis, and chronic anemia [19,20]. In this model, we observed a pronounced erythroid dysregulation, marked by the accumulation of immature erythroblasts and decreased RIV population (Figure 8B). These results highlight the robustness of our spectral cytometry panel in capturing hematopoietic alterations across both age-related and disease-specific contexts, demonstrating its broad applicability for studying hematopoietic heterogeneity.

## 4. Discussion

In this study, we developed and validated a high-dimensional spectral flow cytometry panel capable of simultaneously identifying hematopoietic stem cells (HSC), progenitor subsets, erythroid populations and mature myeloid or lymphoid cells from a single murine bone marrow sample. This method overcomes a major limitation of classical hematopoietic analysis, which typically requires multiple separate panels to profile these different cell populations. By integrating all these analyses into one single assay, we were able to map hematopoietic cells from progenitors to mature cells (Figure 7).

Using this panel, we successfully identified key HSC and progenitor populations, including long-term (LT-HSC) and short-term HSC (ST-HSC), multipotent progenitors (MPP2, MPP3, MPP4), and common myeloid and lymphoid progenitors (CMP, MEP, GMP, CLP) with clear resolution.

Concerning the erythroid compartment, our panel includes the classical markers CD71, CD44 and Ter119. We have chosen to discriminate erythroid sequential maturation stages with the combination of CD71/Ter119 which provided a good discrimination from pro-erythroblasts to reticulocytes [17]. We are aware that others have proposed other strategies such as CD44 vs FSC [4]. However, in our hands and with the cytometer used, the discrimination was not sufficiently clear, perhaps due to weak CD44 signal and size discrimination (data not shown). Furthermore, because OMIQ does not support UMAP analysis that includes a linear parameter such as size discrimination (FSC), inclusion of CD44 would not have allowed to perform unsupervised analysis as proposed here. In the future, improvement of the staining with CD44, or addition of erythroid-specific cell markers may improve erythroid cell annotation.

Last, myeloid and lymphoid cells were sufficiently resolved to identify fractions enriched in Polymorphonuclear cells (PMN, called here granulocyte-enriched), monocytes, macrophages and lymphocytes (B and T cells), enabling general lineage identification and facilitating the gating of lineage-negative populations for HSC and progenitor analysis. The use of additional markers will definitely improve the cell discrimination for both lineages, although one may consider using a lineage-targeted panel to further focus, pending of the results provided first using our approach.

As a proof of validation of the panel, we performed flow cytometry analyses on bone marrow from different mouse models, known to display hematopoietic cell alterations. The panel was applied to anemic disease model, beta-thalassemia mice, demonstrating its ability to capture altered erythropoiesis associated with congenital anemias [19,20]. We also used young and aged animals to capture the related changes in HSC subsets were successfully detected in WT mice, with the expected expansion of long-term HSCs and reduction of multipotent progenitors MPP4 in aged animals compared to young ones [21,22]. These results illustrate that the panel can be reliably used to assess both physiological and pathological hematopoiesis or erythropoiesis, in one shot.

The panel has been designed with the objective to determine the content of bone marrow hematopoietic progenitors and mature cells. Although, other more complex panels have been designated in the literature, most of them are more specialized to study HSPCs, erythroid progenitors, mature myeloid or lymphoid cells. In contrast, here we have chosen to simplify the panel leaving space for additional markers that one would request. In line with this philosophy, we have purposedly left free the FITC/GFP and mCherry spectral signatures since these are often useful for research. Overall, our spectral flow cytometry panel represents a high-resolution tool for preclinical research and for future clinical applications.

## Acknowledgements

This work was supported by funding from the European Union’s Horizon Europe Research and Innovation Programme 2021-2027 under the grant agreement N°101080156 (CL). Funded by the European Union. Views and opinions expressed are however those of the author(s) only and do not necessarily reflect those of the European Union. Neither the European Union nor the granting authority can be held responsible for them. This work was also supported by funding from the Cancéropole PACA (MAL/CF). We thank Hana Hukelova for her administrative support. We thank Jean-Charles Graziano and Arnaud Capel from CRCM’s animal facility. We thank Anne-Laure Bailly from Cytometry platform.

## Author Contributions

LB, CF and MR designed the study and the panel and supervised the experiments and supervised the project and wrote the manuscript. LB, CF and LG performed experiments and analyzed the data. CL, MAL and CF supervised and funded the project. MDG and JG analyzed the data and corrected the manuscript. A.B and J.V corrected the manuscript.

## Supplemental Figure Legends

**Suppl Fig 1. Spectral unmixing and compensation**

The table displays the remaining modifications required after unmixing the acquired data (compensation).

**Suppl Fig 2. PeacoQC – cleaning data**

(A) Percentage of cells that passed (light grey) or did not passed (dark grey) the PeacoQC quality control. (B) The dot plots show each parameter signal during time of acquisition, revealing the presence of cell with altered signal for one or more parameters (not passed). This quality control step may remove up to 30% of the acquired cells, thereby retaining only high-quality cells for downstream analyses.

**Suppl Fig 3. Validation of the expression of cell surface markers on HSC and progenitors**

(A) UMAP as a quality control tool for expression levels of all antibodies in progenitors and HSC are displayed. (B) UMAP as a quality control tool for expression levels of all antibodies only in HSC are displayed. Marker expression scales are shown for each graph and visualized using a rainbow heat scale.

**Suppl Fig 4. Validation of the expression of cell surface markers on erythroid cells**

UMAP as a quality control tool for expression levels of all antibodies in erythroid cells are displayed. Antibodies expression scales are shown in each graph and visualized using a rainbow heat scale.

**Suppl Fig 5. Validation of the expression of cell surface markers on mature myeloid cells**

UMAP as a quality control tool for expression levels of all antibodies in mature myeloid cells are displayed. Antibodies expression scales are shown in each graph and visualized using a rainbow heat scale.

**Suppl Fig 6. Validation of the expression of cell surface markers on all hematopoietic cells**

UMAP as a quality control tool for expression levels of all antibodies in all hematopoietic cells are displayed. Antibodies expression scales are shown in each graph and visualized using a rainbow heat scale.

