## Supplementary figures and images for "Single Spectral Flow Cytometry Panel for Simultaneous Detection of Hematopoietic Stem, Progenitor, and Mature Lineages in Mouse Bone Marrow"

### Supplemental Figure 1

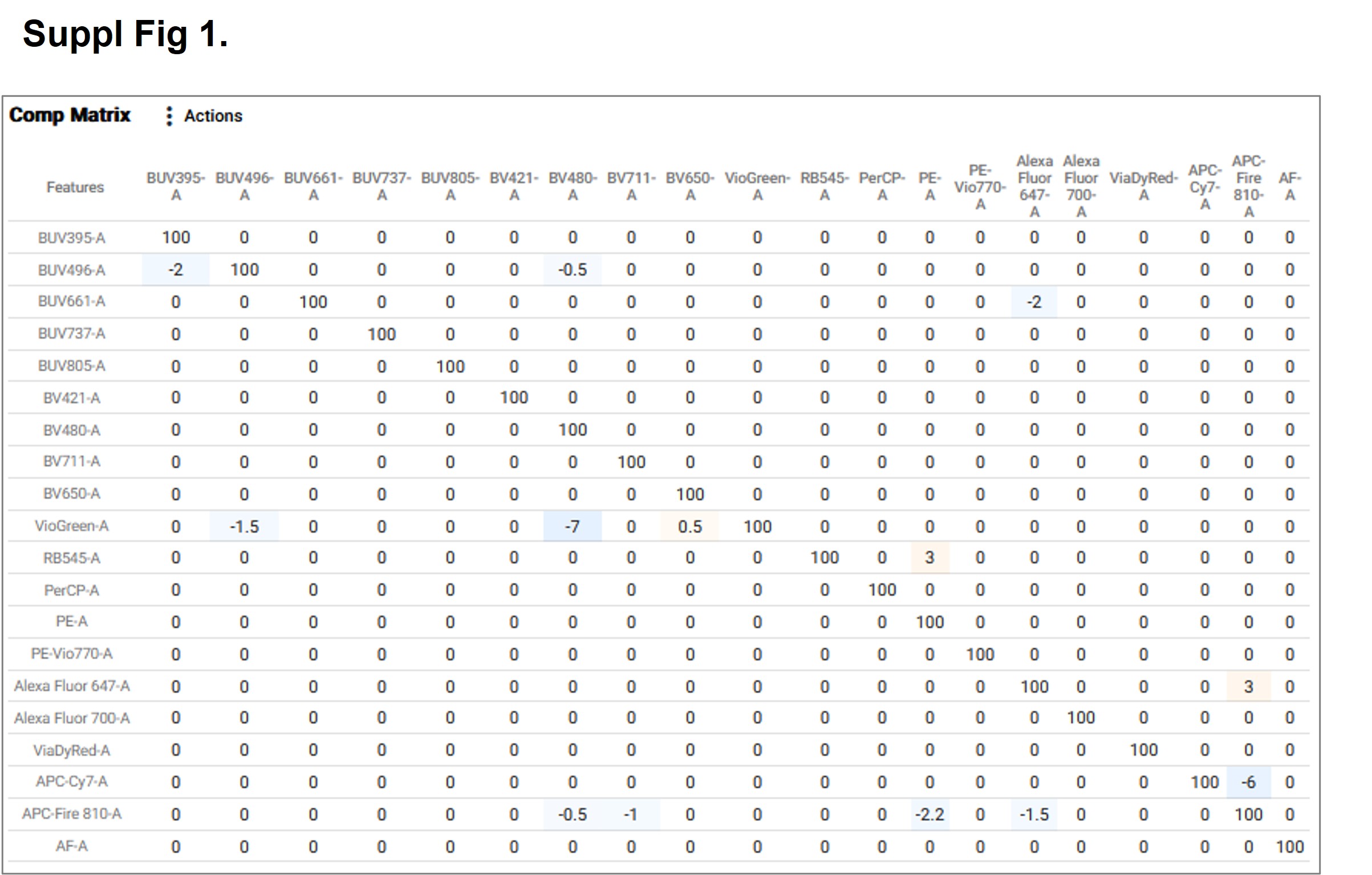

### Supplemental Figure 2

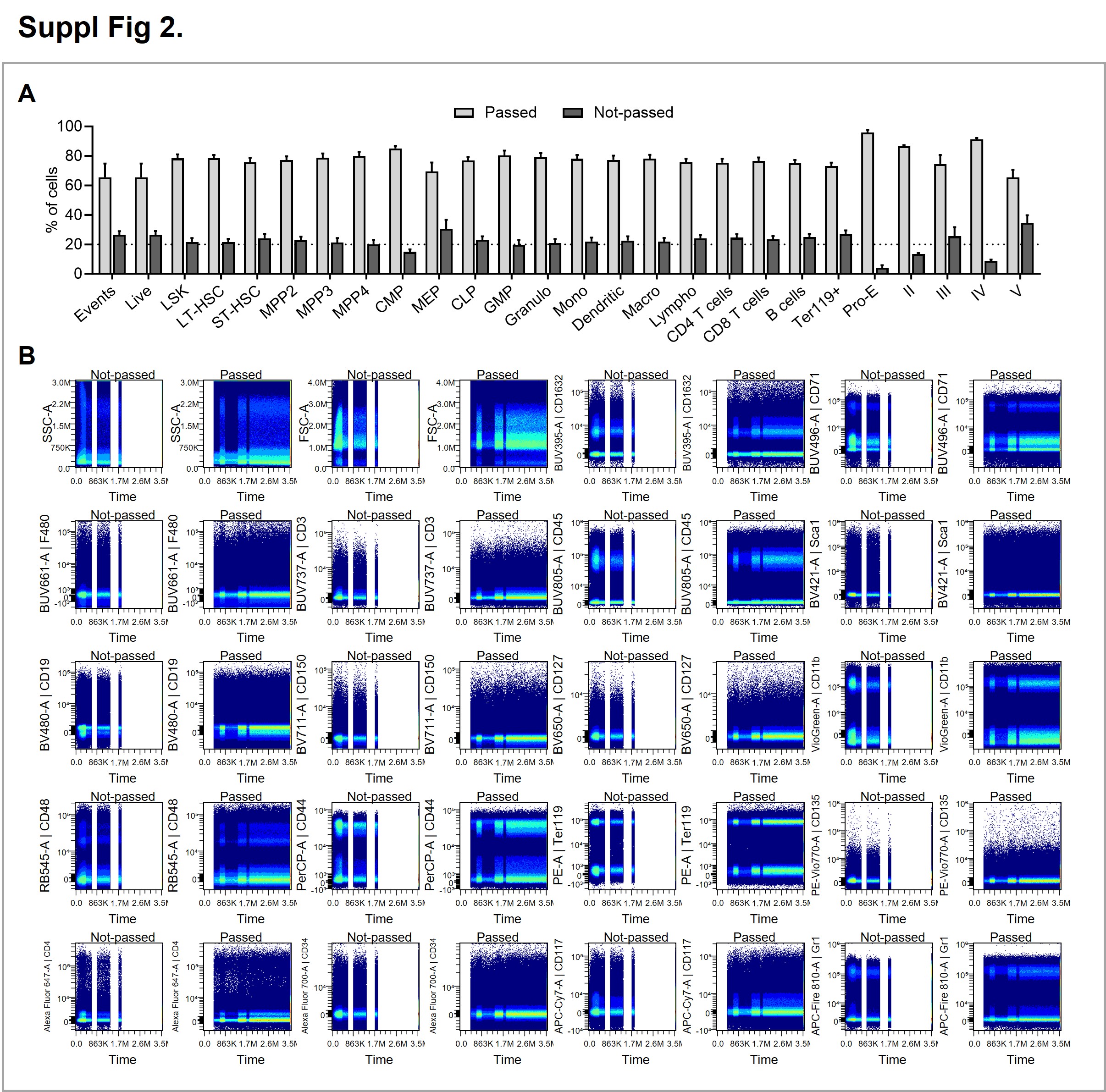

### Supplemental Figure 3

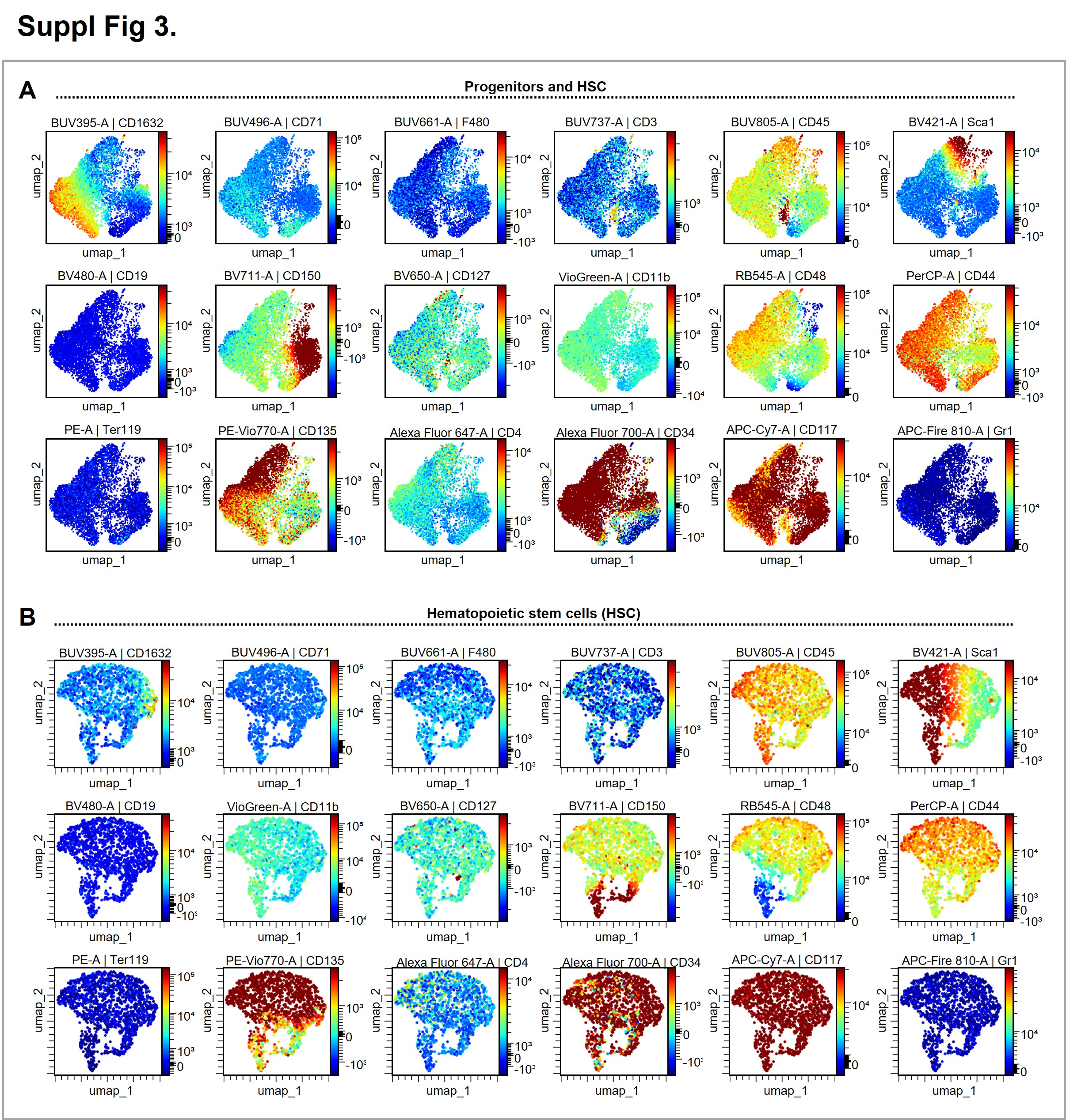

### Supplemental Figure 4

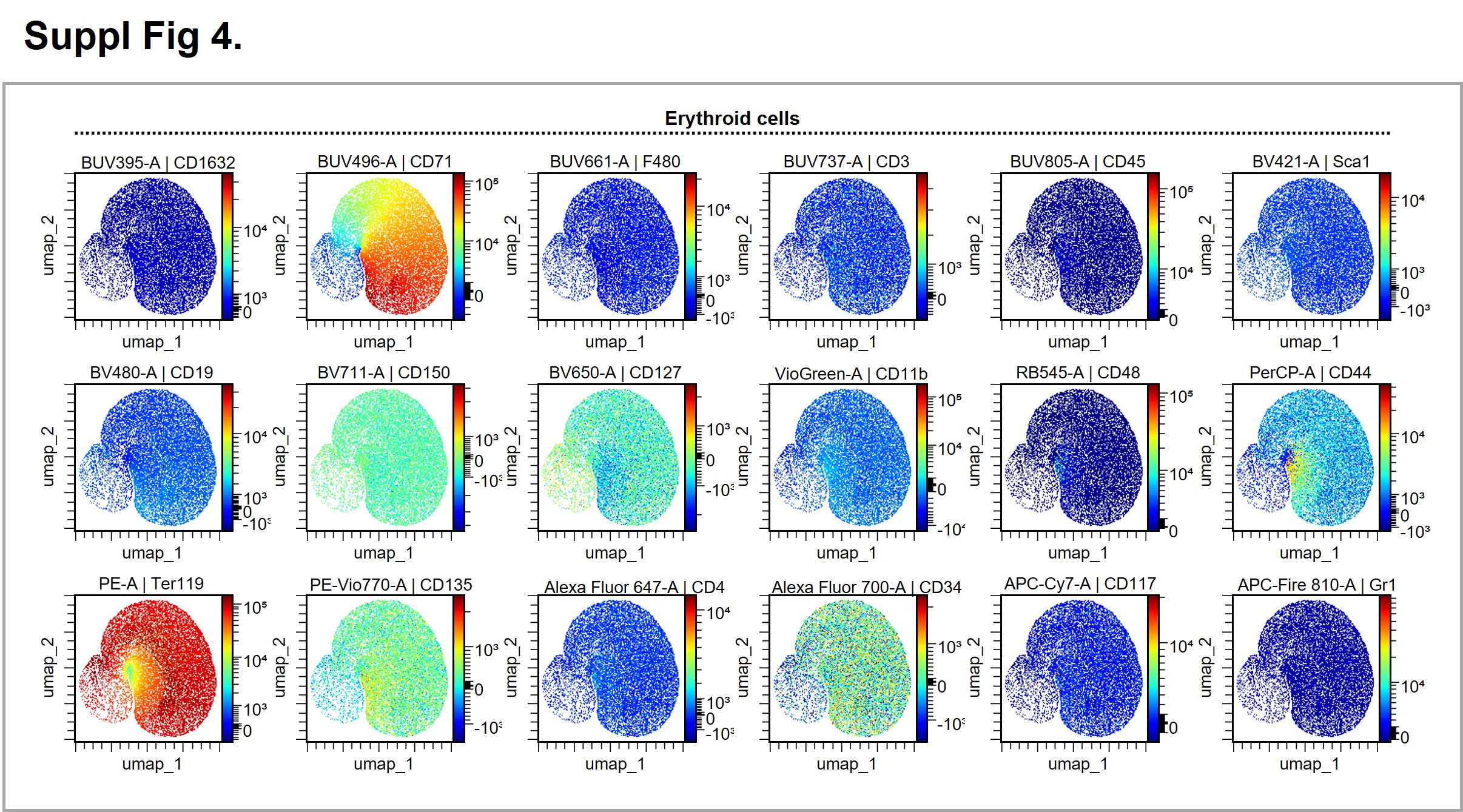

### Supplemental Figure 5

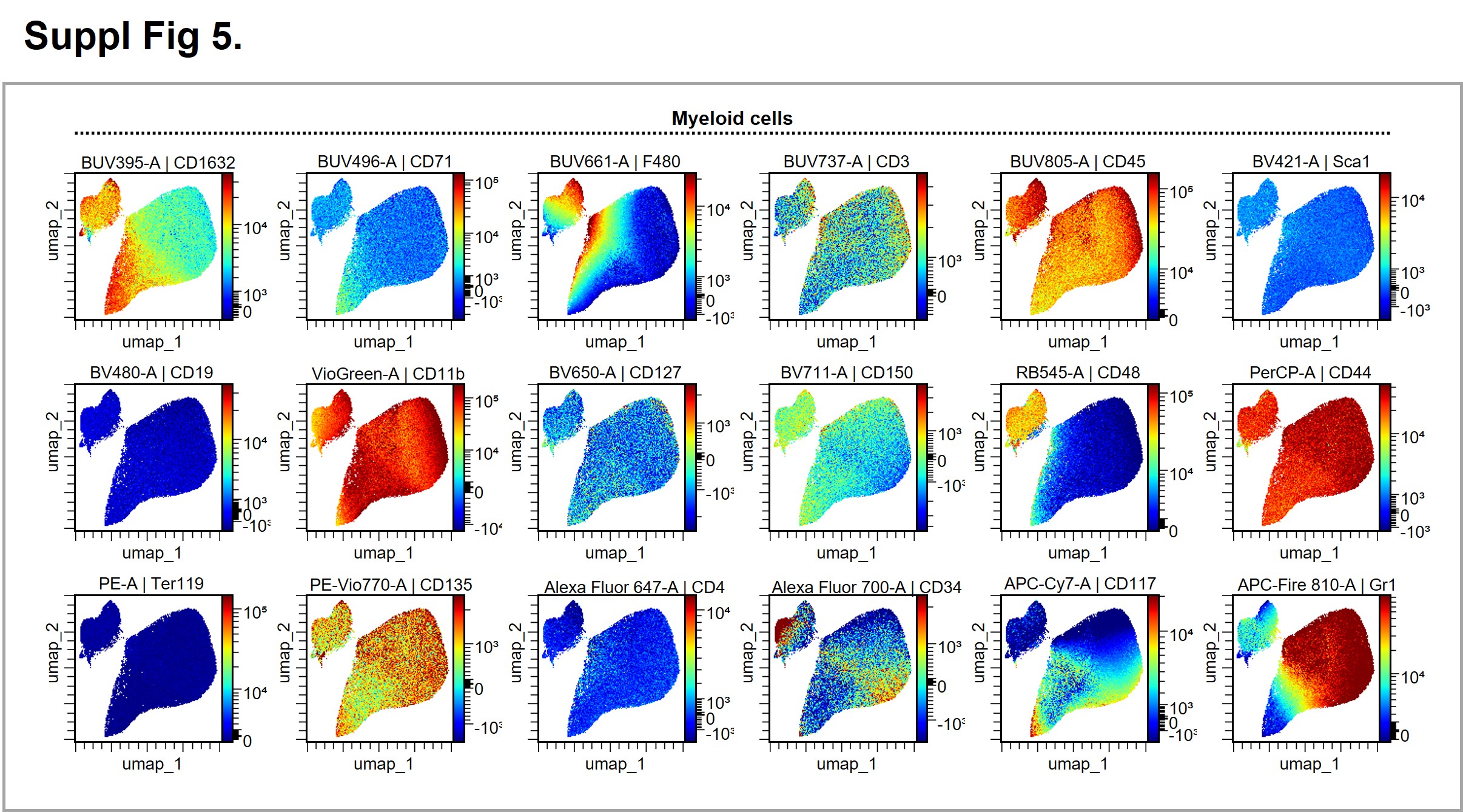

### Supplemental Figure 6

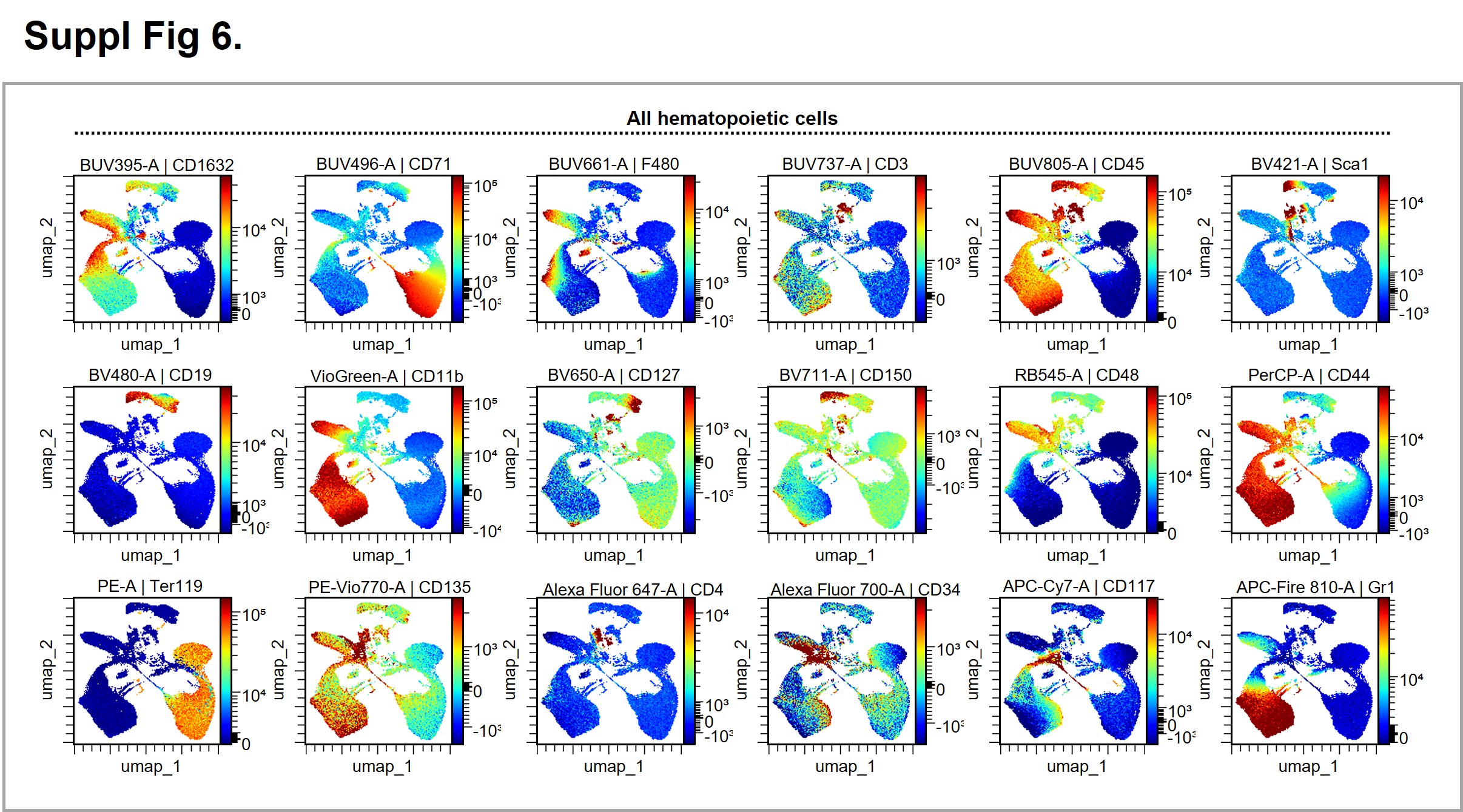
